# Effects of neem extract on *Artemia franciscana*: insights from high-throughput transcriptomics and phenotypic analysis

**DOI:** 10.64898/2026.04.20.719344

**Authors:** Rodolfo Farlora, Paulina Bustos, Eugene M. Tine, Eduardo Jeria, Alex Eapen, Padmakumar Pillai, Jorge Pino, Donald I. Brown, Kathleen Whitlock

## Abstract

Neem-derived biopesticides are increasingly applied in agriculture and have been tested in aquaculture research, yet their effects on non-target aquatic invertebrates remain insufficiently characterized. We evaluated the effect of neem extract on the brine shrimp *Artemia franciscana* using an integrated ecotoxicological approach combining phenotypic, transcriptomic, and histological analyses. Juvenile *A. franciscana* exhibited dose-dependent mortality and sublethal abnormalities, with a 24 h median lethal concentration of 292.48 mg/L (95% confidence interval, 257.75– 331.89) for mortality and a median effective concentration of 146.36 mg/L (95% confidence interval, 113.04–189.50) for the combined endpoint “abnormal + dead”. In adults, males showed greater mortality than females after extended exposure. High-throughput RNA sequencing revealed broad treatment-associated differences in transcript abundance, with juveniles displaying downregulation of detoxification enzymes and chitin biosynthesis genes, alongside enrichment of immune- and cuticle-related gene ontologies. Adults showed transcriptional signatures of stress, including upregulation of heat shock proteins and cytoskeletal components, and suppression of genes involved in energy metabolism. Chitin precursor enzymes were selectively downregulated in males, and altered carbohydrate metabolism was observed in females. Histological analyses revealed structural deterioration of the brood sac cuticle and reduced ovarian area in treated females, consistent with transcriptomic evidence of impaired exoskeletal and reproductive processes. Overall, neem exposure was associated with phenotypic, histological, and transcriptomic changes in *A. franciscana*. These results support the use of combined transcriptomic and histopathological endpoints to characterize responses to plant-derived biopesticides in aquatic arthropods.

## 1. Introduction

The neem tree (*Azadirachta indica*) produces bioactive compounds, primarily azadirachtins, widely employed as botanical insecticides (1,2). Neem-derived formulations have been increasingly adopted in agriculture, and have been tested experimentally in aquaculture settings, where studies have reported immunostimulant and antiparasitic effects in fish (3–6). However, the use of neem-based products in systems connected to natural or semi-natural water bodies raises concerns about potential adverse impacts on non-target aquatic organisms, particularly invertebrates that underpin food-web structure (7).

Transcriptomics has emerged as a valuable tool in ecotoxicological studies of aquatic species, offering insights into the molecular mechanisms underlying toxicant exposure and facilitating the development of sensitive molecular biomarkers (8–10). High-throughput transcriptome sequencing (HTS) can provide broad surveys of gene expression changes associated with toxicant exposure, allowing the identification of candidate responsive genes and biological pathways (11,12). Increasingly, such mechanistic information is organized within Adverse Outcome Pathway (AOP) frameworks, which link molecular and cellular key events to organism- and population-level adverse outcomes for use in ecotoxicological risk assessment (13,14).

The brine shrimp, *Artemia franciscana*, serves as a valuable model for comparative toxicology and invertebrate physiology, combining ease of culture and sensitivity to chemically induced physiological disruption (15–18). Although acute toxicity and general physiological responses to various contaminants in the genus *Artemia* have been extensively investigated (19,20), mechanistic data describing how neem-derived compounds affect molecular pathways and tissues in this branchiopod remain limited, despite the growing use of azadirachtin-based neem biopesticides and evidence of adverse effects in other aquatic invertebrates (7,21). To our knowledge, no previous study has combined transcriptomic and histological endpoints to investigate sublethal effects of neem exposure in *A. franciscana*.

To address this gap, we combined traditional phenotypic endpoints (survival and developmental abnormalities), RNA-seq transcriptomics and histological examinations to evaluate neem extract toxicity in juvenile and adult *A. franciscana*. Specifically, we (i) quantified dose-dependent mortality and developmental abnormalities in juveniles, (ii) characterized adult survival patterns and reproductive condition under subchronic exposure, (iii) identified life stage- and sex-specific transcriptomic responses to neem treatment, with a particular focus on chitin biosynthesis and cuticle-related pathways, and (iv) assessed histological alterations in adult reproductive tissues. Together, these data provide an integrated assessment of phenotypic, histological, and transcriptomic responses to neem exposure in this model crustacean, and identify candidate biological processes associated with treatment.

## 2. Materials and Methods

### 2.1. Preparation of neem extract

Neem kernel extract stock solution (3 g/L) was prepared from a commercial product (Coromandel/Murugappa; Parry’s mBio NeemAzal® Technical Pack No 06/1/3, Lot No B365). This formulation contains azadirachtin A (25–50%) along with other naturally occurring kernel constituents, including additional azadirachtins (14–21%), limonoids (1–10%), and fatty acids (1–2%). Because of their low solubility in water, dilutions were made in 0.5% dimethyl sulfoxide (DMSO) in artificial seawater. The control group received artificial seawater containing 0.5% DMSO only, corresponding to the solvent concentration used in all neem extract dilutions. The stock was stirred overnight at room temperature before use. Neem concentrations are reported as ppm (mg/L).

### 2.2. Culture of juvenile and adult Artemia

*A. franciscana* were maintained within a salinity range of 35–40 ppt at 28 °C on a controlled light-dark cycle of 14 h light/10 h dark. Juvenile Artemia were collected 24 hours after initiation of the cultures (nauplii stages: 12-24 hours post culture) and placed in 24 well plates with approximately 50 juveniles per well. Artificial sea water was removed, and 2 mL of test solutions were added and allowed to incubate for 24 hours. The results were expressed in terms of Lethal Concentration (LC). The Effective Concentration (EC), representing the percentage of abnormal and deceased animals within the population, was also calculated.

Adult experimental groups were maintained in six-well culture plates, with three individuals of the same sex per well. Each concentration was tested in six replicate wells (n = 18 adults per sex and treatment). Animals were exposed to neem extract for seven days, with daily renewal of test solutions and feeding according to the standard culture regime. Because no consistent sublethal abnormalities could be reliably scored in adults, mortality was recorded daily throughout the 7-day exposure period.

Overall, the experimental design comprised an acute 24 h juvenile exposure (0–900 ppm) used for toxicity assessment and juvenile RNA-seq in the sublethal range (0–400 ppm), and a 7-day adult exposure that supported three endpoints: survival analysis at 0, 250 and 500 ppm, adult RNA-seq at 0, 250 and 500 ppm, and histology assessments at 0, 100, 250 and 500 ppm.

### 2.3. Total RNA extraction and construction of cDNA libraries

After the 24-hour exposure to the sublethal range of neem extract (0 ppm control, 100, 200, 300 and 400 ppm), with each treatment performed in triplicate (three wells per concentration; approximately 50 juveniles per well), juveniles were collected from each well using filter paper, preserved in RNAlater (Thermo Fisher Scientific), and stored at −80 °C. Each well was treated as one biological replicate, yielding three independent RNA samples per juvenile treatment.

Adult male and female *Artemia* were exposed for seven days to neem extract at 0 ppm (control), 250 ppm, and 500 ppm. For each sex and concentration, three RNA-seq libraries were prepared. Each library consisted of a pool of three surviving adults collected at day 7 from the same sex and treatment group. When necessary, individuals were combined from more than one well within the same sex and treatment group to obtain the number required for library construction.

Total RNA was extracted from pooled brine shrimp using the TRIzol reagent (Invitrogen) protocol. RNA purity was assessed using the A260/A280 ratio with a NanoDrop Lite spectrophotometer (Thermo Fisher Scientific), and RNA integrity was verified by visualizing samples in agarose gels under denaturing conditions. Subsequently, a total of 33 double-stranded cDNA libraries were generated using the TruSeq® RNA Sample Preparation Kit v2 (Illumina®, USA) by Novogene USA, corresponding to three biological replicates per treatment and life stage/sex combination (five juvenile treatments × three replicates, and three adult treatments × two sexes × three replicates). All libraries were sequenced on the NovaSeq 6000 platform (Illumina®) with 2 × 150 bp paired-end reads. The list of sequenced cDNA libraries is detailed in **S1 Table**. The original sequencing files have been uploaded to NCBI’s Sequence Read Archive (SRA) under the BioProject ID PRJNA1402122.

### 2.4. Sequencing of cDNA libraries and bioinformatics analysis

Transcriptomic analyses were conducted as an exploratory assessment of treatment-associated transcript abundance patterns using a *de novo* assembled reference transcriptome. Following the removal of adapters and low-quality sequences, the clean sequences were individually trimmed and *de novo* assembled into a unified file using CLC Genomics Workbench software (Version 11.0.1, CLC Bio, Denmark). The assembly parameters included a mismatch cost of 2, an insert cost of 3, a minimum contig length of 400 bp, a similarity threshold of 0.9, and a trimming quality score of 0.05. This assembly yielded 88,367 contigs (S1 Dataset) that were subsequently annotated to the NCBI non-redundant and UniProtKB/SwissProt Protein databases using the BLASTx tool (S2 Dataset).

To identify over and under-represented Gene Ontology (GO) categories related to molecular function, biological processes, and cellular components, we conducted GO functional enrichment analysis using the Rank-based Gene Ontology Analysis with Adaptive Clustering (RBGOA) tool (22). Log_2_ Fold Change (log2FC) and p-value were employed to evaluate the strength and significance of gene expression (23). Only sequences with an expected value threshold (E-value) of similarity to sequences in the UniProt database less than 0.01 were considered for analysis.

The annotated transcriptome served as a reference for differential gene expression (DEGs) analysis by RNA-seq. Clean sequences obtained from each library were aligned to the reference transcriptome using CLC Genomics Workbench software. Expression values were calculated as transcripts per million mapped reads (TPM) (S3 Dataset; S4 Dataset). DEGs, meeting the criteria of |log2FC| ≥ 4 and a false discovery rate (FDR) p-value < 0.05 were identified and visualized in hierarchical clustering heat maps. Hierarchical clustering employed Euclidean distance and full linkage as parameters for analysis.

### 2.5. In silico identification of genes associated with chitin synthesis

To investigate chitin synthesis in *A. franciscana*, a gene panel was curated from the assembled transcriptome, selecting contigs associated with this process through bioinformatic analysis and literature search on the genes and pathways associated with chitin metabolism in various crustacean species (24). Homology searches against the NR protein database (E-value < 0.005) identified sequences related to chitin synthesis. These gene panels were used as reference datasets and reads from larvae and adults across treatment replicates were mapped against them using CLC Genomics Workbench, applying the same RNA-seq settings and hierarchical clustering parameters as described previously. Expression levels for the selected genes were quantified as transcripts per million (TPM) derived from the mapped libraries, and these values were used for subsequent analyses.

### 2.6. Histological analysis of adult A. franciscana

A total of 80 adult *A. franciscana* were selected for histological evaluation (10 males and 10 females per concentration), and these individuals were reserved exclusively for histological processing. Animals were fixed in Bouin’s solution for 48 hours and subsequently rinsed under running tap water. Standard histological procedures were followed, including dehydration through a graded ethanol series, clearing in butanol, and embedding in Paraplast Plus® (Merck). Embedded tissues were sectioned at 5 μm thickness using a Leica RM2255 microtome and mounted on glass slides. For visualization, tissue sections were dewaxed and rehydrated via a xylene-butanol sequence and processed for trichrome staining: Hematoxylin (Merck) for nuclei, Erythrosin-orange G (Merck) for cytoplasmic differentiation, and Aniline blue (Merck) for connective tissues and secretion granules. Slides were dehydrated in ethanol, cleared in xylol (Merck), and mounted in Entellan™ (Merck). A Leitz-Leica DMRBE microscope equipped with a Leica DFC290 digital camera was used to document histological features.

The gonadal areas of male and female individuals were identified and photographed, with images taken from at least three consecutive sections per individual. Gonadal area measurements were performed from digital micrographs using ImageJ software (version 1.54h; National Institutes of Health) after calibration with the microscope scale bar. Areas were quantified in at least three consecutive sections per individual and averaged for downstream analysis. In mature females, additional measurements focused on regions corresponding to embryos in the brood sac (EBS). Representative histological images, including EBS deformities in treated females, are presented in the main figures, and additional quantitative data on gonadal area measurements are provided in the supplementary material.

### 2.7. Statistical analysis

Statistical analyses were conducted to evaluate the effects of neem extract exposure on *A. franciscana* at phenotypic, transcriptomic, and histological levels. Phenotypic data, including mortality and developmental abnormalities in juveniles, were analyzed using one-way analysis of variance (ANOVA), followed by Tukey’s multiple comparison test to determine significant differences between groups (*P* < 0.05). For adult survival analysis, a Kaplan-Meier survival plot was generated using GraphPad Prism (10.4.1), with statistical differences evaluated using the log-rank (Mantel-Cox) test in the same software.

For transcriptomic analysis, Principal Component Analysis (PCA) was performed on transcripts per million (TPM) values derived from paired-end cDNA libraries, projecting samples onto a two-dimensional space defined by the first and second principal components of the covariance matrix. Differential gene expression analysis was conducted using a T-test with false discovery rate (FDR) correction for multiple comparisons. Independent filtering was applied using a cutoff of |log2FC| ≥ 4 to minimize false positives, and a final subset of transcripts was retained for further analysis, with statistical significance set at *P* < 0.05. All RNA-seq statistical analyses were performed using CLC Genomics Workbench software.

For the expression analysis of chitin-related genes, TPM values were used as input for statistical comparisons, and gene expression differences in juveniles were assessed using one-way ANOVA, followed by Tukey’s multiple comparison test (*P* < 0.05). In adults, chitin-related gene expression was analyzed using two-way ANOVA, followed by Tukey’s multiple comparison test to evaluate differences between treatment groups (*P* < 0.05).

For histometric data, normality was assessed using the Shapiro–Wilk test. In cases where data did not meet normality assumptions, Box-Cox transformation was applied (25). Statistical comparisons between experimental conditions were performed using one-way ANOVA, followed by Tukey’s multiple comparison test (*P* < 0.05).

All statistical analyses for phenotypic data, chitin-related gene expression, and histometric data were conducted using GraphPad Prism software (10.4.1).

### 2.8 Ethics statement

This study involved *Artemia franciscana*, a non-cephalopod invertebrate, and did not involve human participants, vertebrate animals, cephalopods, or field research. Therefore, institutional ethics approval was not required.

### 2.9. Artificial intelligence tools and technologies

During manuscript preparation, the authors used ChatGPT (OpenAI) to assist with language editing and to improve the clarity and consistency of the manuscript text. All AI-assisted output was reviewed and edited by the authors for accuracy and consistency with the original results and interpretations. No primary data, analyses, results, figures, tables, or Supporting Information files were generated by the AI tool. The authors take full responsibility for the content of the manuscript.

## 3. Results

### 3.1. Neem extract induces dose-dependent mortality and abnormal phenotypes in juvenile A. franciscana

To assess the effects of neem extract on juvenile *A. franciscana*, nauplii were exposed to concentrations ranging from 0 to 900 ppm and scored after 24 h. Individuals showing active swimming with coordinated undulating movements of their podia were classified as normal (S1 Video). Those showing reduced and/or uncoordinated podial movements and an impaired ability to remain in the water column were classified as abnormal. Individuals that remained immobile at the bottom of the well, showed no response to gentle tapping, and exhibited necrotic morphology were classified as dead (S1 Video). At 24 h, all juveniles in the control group (0 ppm) exhibited a normal phenotype (**Fig 1A**). At concentrations below 100 ppm, most juveniles remained normal. Between 100 and 300 ppm, the proportion of abnormal juveniles increased significantly (P < 0.001), accompanied by a marked reduction in the proportion of normal individuals. At 200 ppm, only ∼17% of juveniles remained normal, whereas abnormal and dead individuals accounted for the majority of the population. The proportion of abnormal juveniles peaked at 300 ppm (∼64%) and then declined at higher concentrations as mortality became the predominant response. Mortality increased markedly from 400 ppm onward, exceeded 85% at 600 ppm, and reached 100% at 900 ppm. Based on these concentration-response data, the 24 h LC₅₀ for juvenile mortality was 292.48 ppm (95% CI: 257.75–331.89), and the EC₅₀ for the combined endpoint “abnormal + dead” was 146.36 ppm.

**Fig 1.**
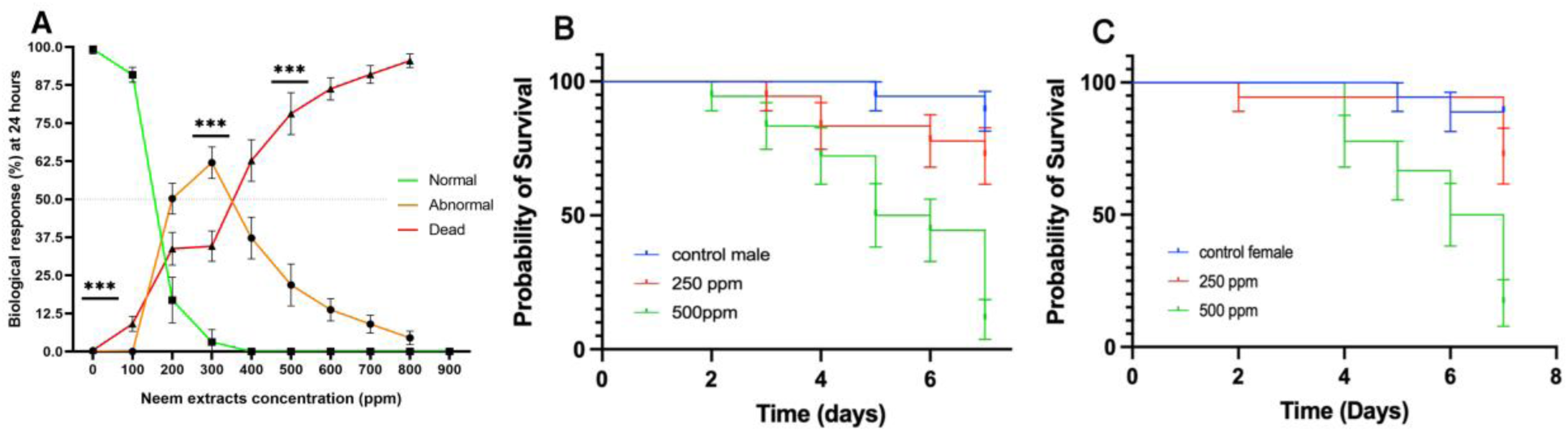
Dose-dependent effects of neem extract on survival and developmental phenotype of *Artemia franciscana*. **A**) Juveniles exposed for 24 h. The proportion of individuals displaying a normal phenotype (green) decreased with increasing neem concentration, while abnormal phenotypes (orange) increased, peaking at 300 ppm, and declined at higher concentrations as mortality (red) increased. Data are shown as mean ± SD. Statistical differences were evaluated using one-way ANOVA followed by Tukey’s multiple comparison tests. **(B, C)** Survival of adult *Artemia*. Kaplan–Meier survival curves are shown for (B) males and (C) females exposed to 0, 250, and 500 ppm neem. Statistical differences were evaluated using the log-rank (Mantel–Cox) test.

### 3.2. Neem extract exposure reduces adult survival in A. franciscana

Preliminary observations indicated that adult *A. franciscana* were more resistant to neem extract than juveniles, which exhibited phenotypic effects within 24 hours of exposure. Therefore, the potential effects of neem extract on adults were assessed over a period of 7 days. Males and females were analyzed separately, with males identified by the presence of claspers, and only gravid females included in the study (see **S2 Video**). At the lower concentration tested (250 ppm), some mortality was observed by the end of the 7-day exposure, but the difference from controls was not statistically significant. After seven days of exposure to 500 ppm, both males and females showed pronounced mortality (>80% in each sex; **Fig 1B** for males, **Fig 1C** for females), whereas control groups exhibited low mortality (approximately 11% by day 7, with deaths beginning on day 5). At both 250 and 500 ppm, male survival declined earlier than female survival in the plotted trajectories. During the exposure period, females in all groups continued to release free-swimming nauplii (ovoviviparous reproduction).

### 3.3. RNA-seq reveals widespread gene expression changes in neem-treated A. franciscana

To explore transcript-level responses associated with neem exposure, we performed a *de novo* high-throughput transcriptome assembly to serve as a reference for differential expression analysis. After quality filtering, the sequencing yielded 1,604,747,440 clean reads (**Table 1**), which assembled into 88,367 contigs with an N50 of 1,120 bp. About 70% of contigs (61,999 sequences) were successfully annotated against protein databases, which served as the reference for differential expression analysis. Across all samples, the exploratory screening identified a broad set of candidate treatment- responsive transcripts. Principal component analysis (PCA) of the global gene expression profiles (**Fig 2**) revealed a clear separation between juveniles and adults along the first principal component (**Fig 2A**). When analyzed within each life stage, distinct clustering of neem-treated vs. untreated (control) transcriptomes was observed for both juveniles and adults. In juveniles, all neem-treated groups separated from controls along PC1 (**Fig 2B**). In adults, transcriptional profiles of treated individuals also diverged from controls (**Fig 2C**). Additionally, the second principal component (PC2) distinctly separated adult males from adult females, reflecting underlying sex-specific transcriptional differences in Artemia. Notably, the number of neem-responsive genes was much higher in adults (19,957 DEGs) than in juveniles (933 DEGs).

**Fig 2.**
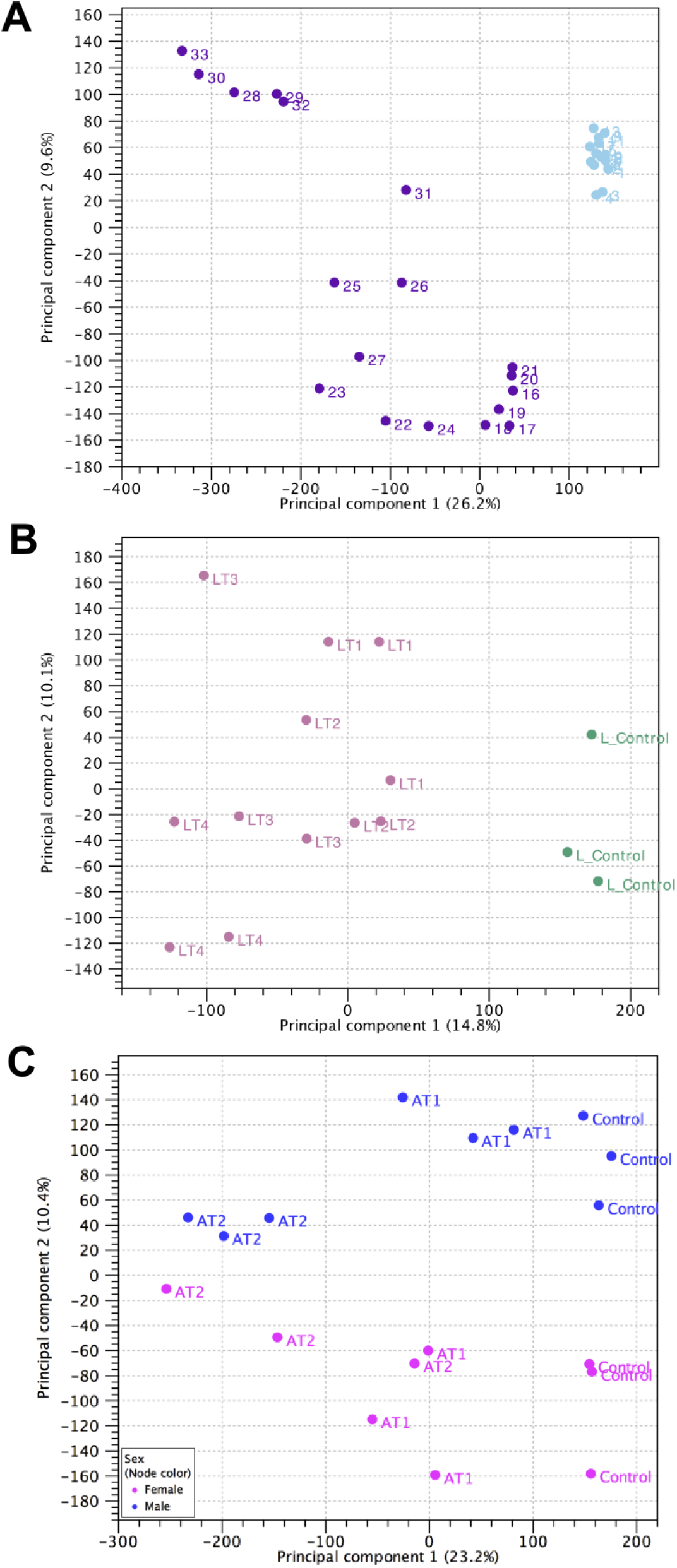
Principal component analysis (PCA) of global gene expression profiles in *A. franciscana* exposed to neem extracts. **(A)** PCA of all RNA-seq libraries (n = 33) showing separation between juveniles and adults along the first principal component (PC1). Each point represents one library; colours and/or symbols indicate life stage, sex and treatment (see panel legend). **(B)** PCA of juvenile libraries only (0, 100, 200, 300 and 400 ppm of neem extract), showing separation between control and neem-treated groups along PC1. **(C)** PCA of adult libraries only (0, 250 and 500 ppm of neem extract), illustrating differences between control and treated samples and separation between males and females.

**Table 1:**
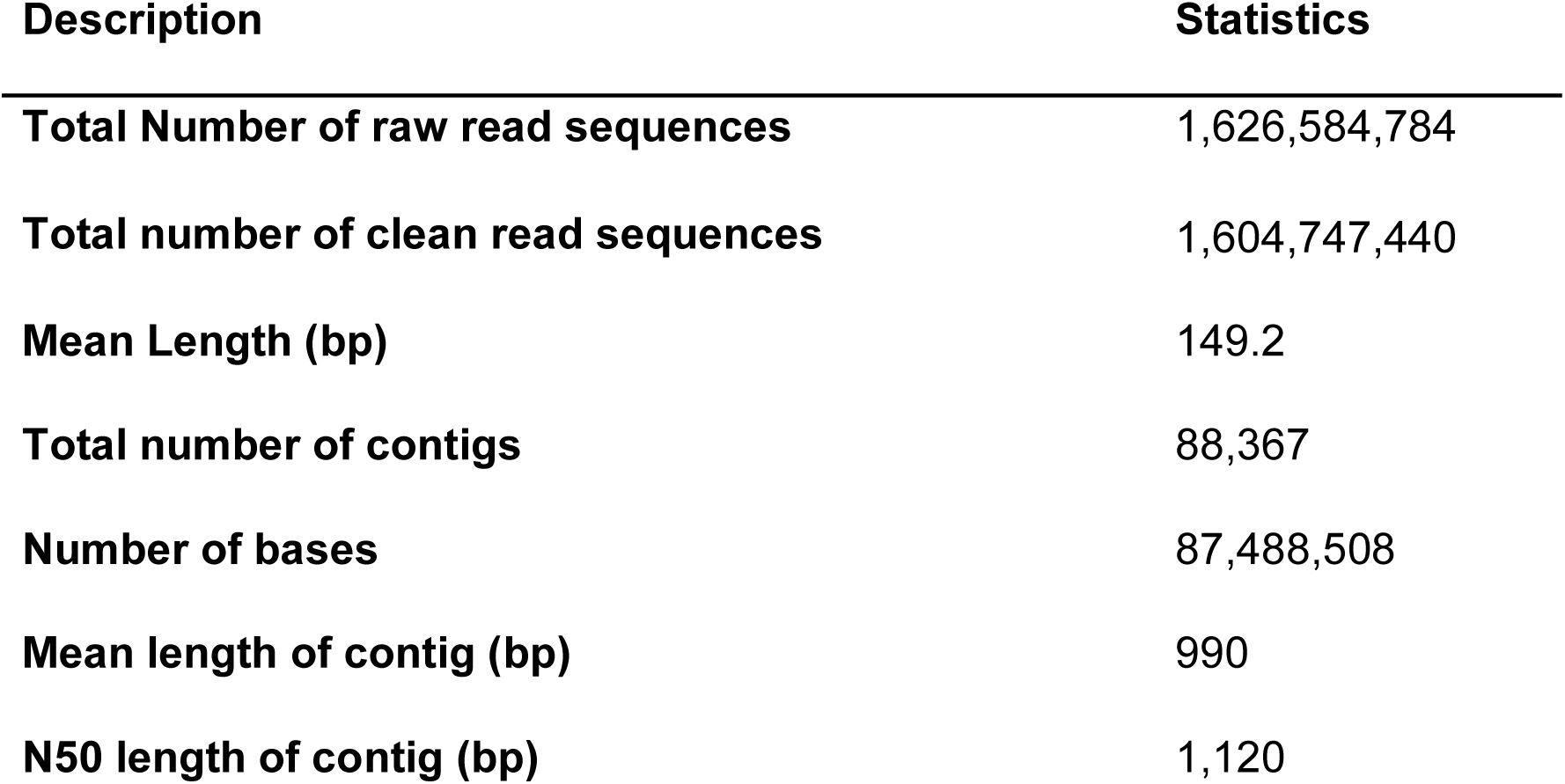
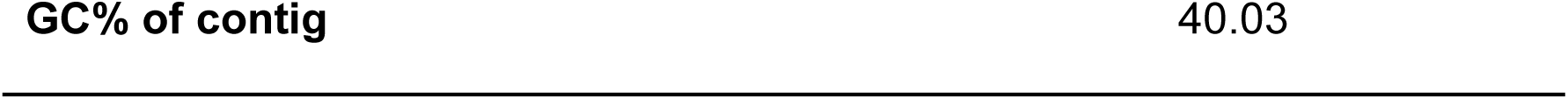
Summary of RNA-seq output and de novo transcriptome assembly statistics for *Artemia franciscana*. Values are based on 33 cDNA libraries from juveniles and adults and include total raw and clean reads, mean read length, number of contigs, total assembled bases, mean contig length, N50 and GC content.

| Description | Statistics |
| --- | --- |
| Total Number of raw read sequences | 1,626,584,784 |
| Total number of clean read sequences | 1,604,747,440 |
| Mean Length (bp) | 149.2 |
| Total number of contigs | 88,367 |
| Number of bases | 87,488,508 |
| Mean length of contig (bp) | 990 |
| N50 length of contig (bp) | 1,120 |

### 3.4. Transcript abundance patterns in juvenile A. franciscana exposed to neem are consistent with metabolic and cuticle-associated responses

Hierarchical clustering of differentially expressed genes (DEGs) revealed distinct transcriptional differences between untreated and neem-treated juvenile *A. franciscana*, with control samples forming a separate cluster from all exposed groups (**Fig 3A**). Among neem-treated juveniles, the 100 and 200 ppm groups clustered together, whereas the 300 and 400 ppm groups showed partially overlapping but distinct profiles.

**Fig 3.**
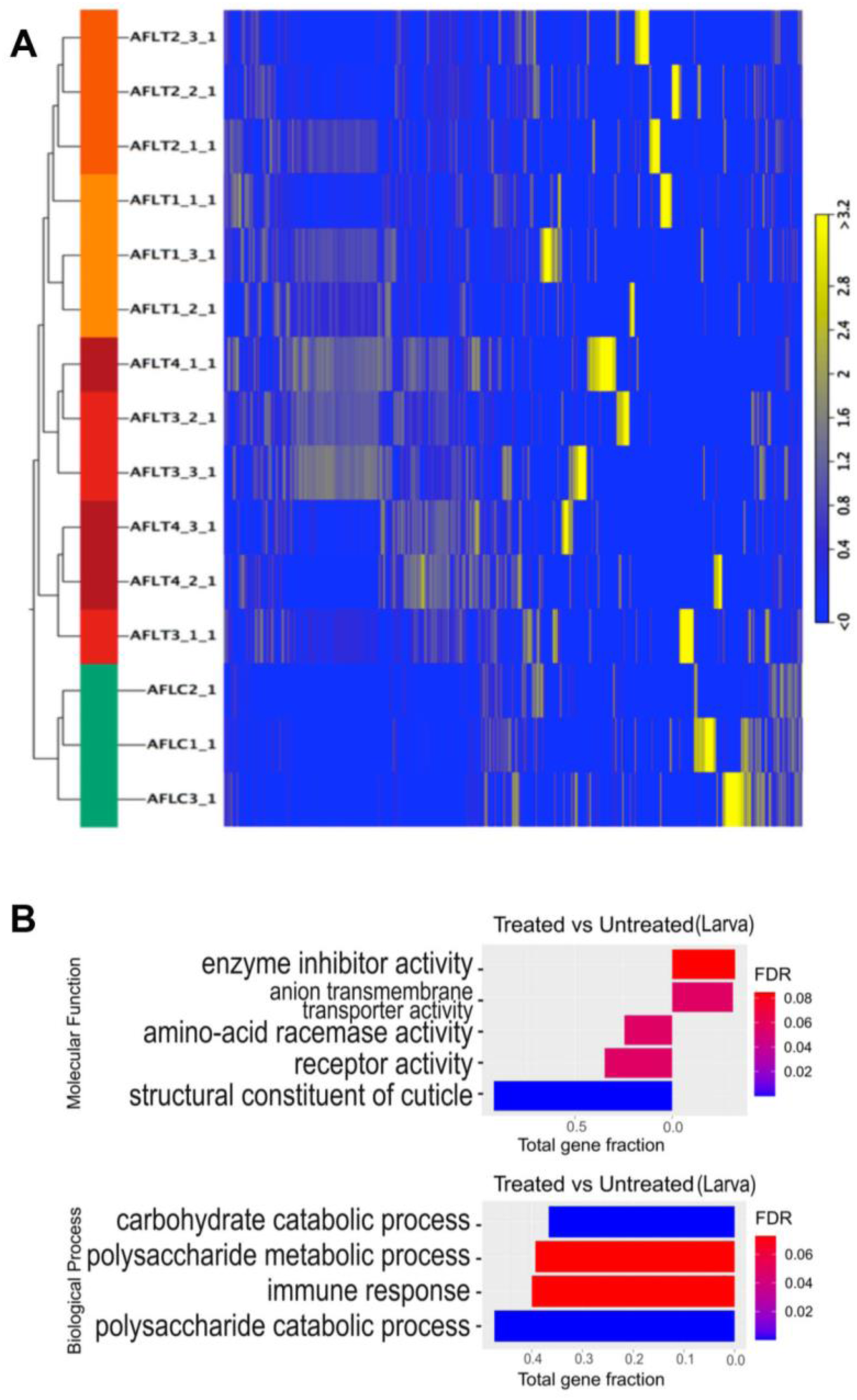
Transcriptomic responses to neem extract exposure in juvenile *A. franciscana*. **(A)** Heat map of hierarchical clustering of differentially expressed genes (DEGs) from juvenile *A. franciscana* exposed for 24 h to 0 (control) or 100, 200, 300 and 400 ppm neem (n = 3 libraries per treatment; 15 libraries in total). Columns represent individual libraries (controls in green; neem-treated groups in progressively darker warm colours from 100 to 400 ppm). The colour scale indicates relative transcript abundance, from lower (blue) to higher (yellow) expression. **(B)** Enriched GO categories for molecular function in treated (left) and untreated (right) juveniles. Bar length indicates the number of DEGs assigned to each GO term. “Total gene fraction” denotes the proportion of all annotated genes in the transcriptome associated with that term. Only terms with P < 0.01 are shown. The colour gradient reflects false discovery rate (FDR), with blue indicating lower FDR (higher confidence) and red indicating higher FDR.

Differential expression analysis (**Table 2**) identified a set of genes with pronounced changes in expression following neem exposure. Strongly induced genes included *pseudouridine-5′-phosphatase* (log₂FC = 3.04), an enzyme involved in RNA modification, and *GTP cyclohydrolase 1* (log₂FC = 2.99), which participates in folate biosynthesis. Other upregulated transcripts included *cathepsin S* (log₂FC = 2.97), a lysosomal protease associated with protein catabolism, and *mitochondrial uncoupling protein 3* (log₂FC = 2.87), a protein linked to regulation of mitochondrial membrane potential.

**Table 2.**
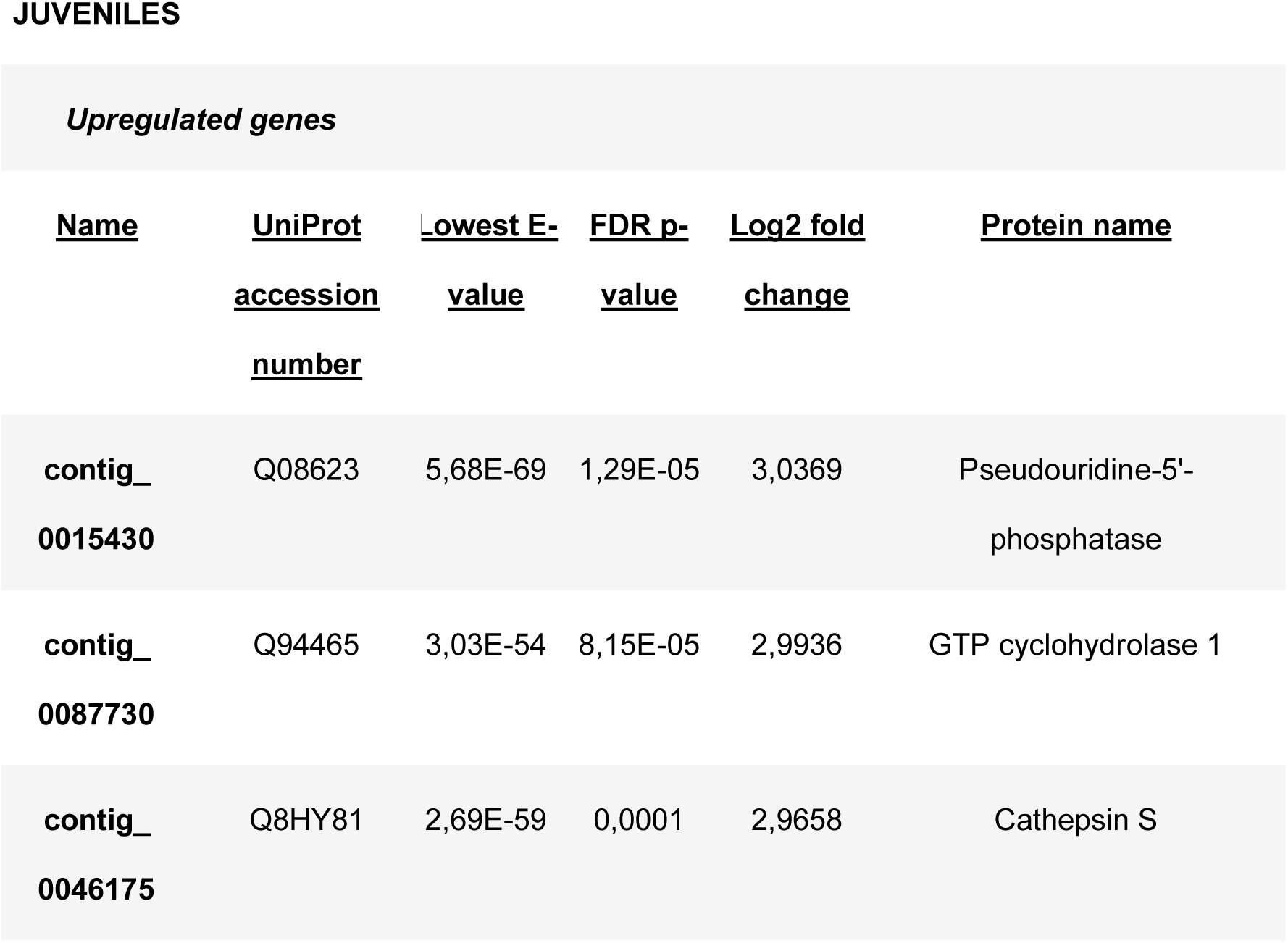
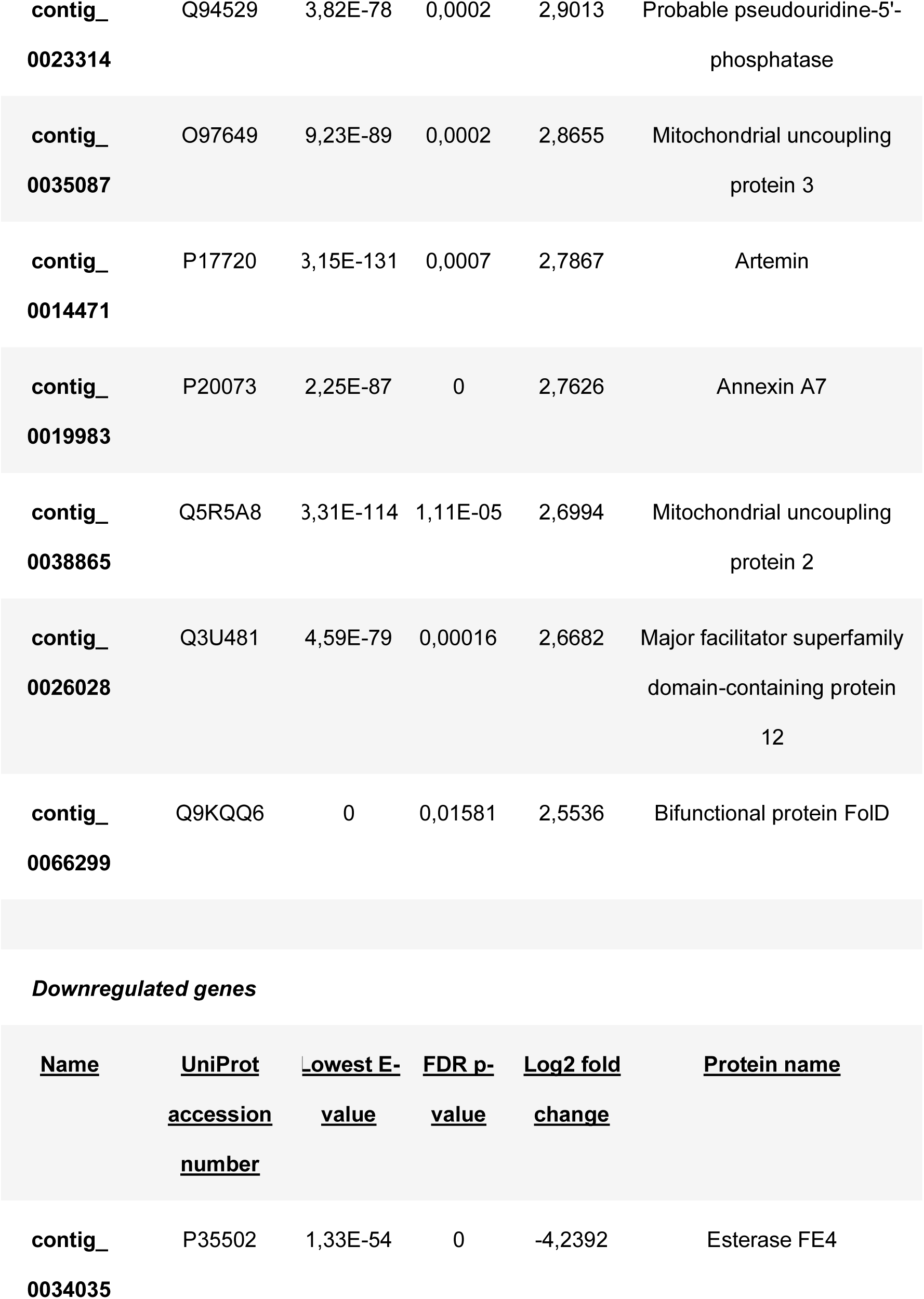
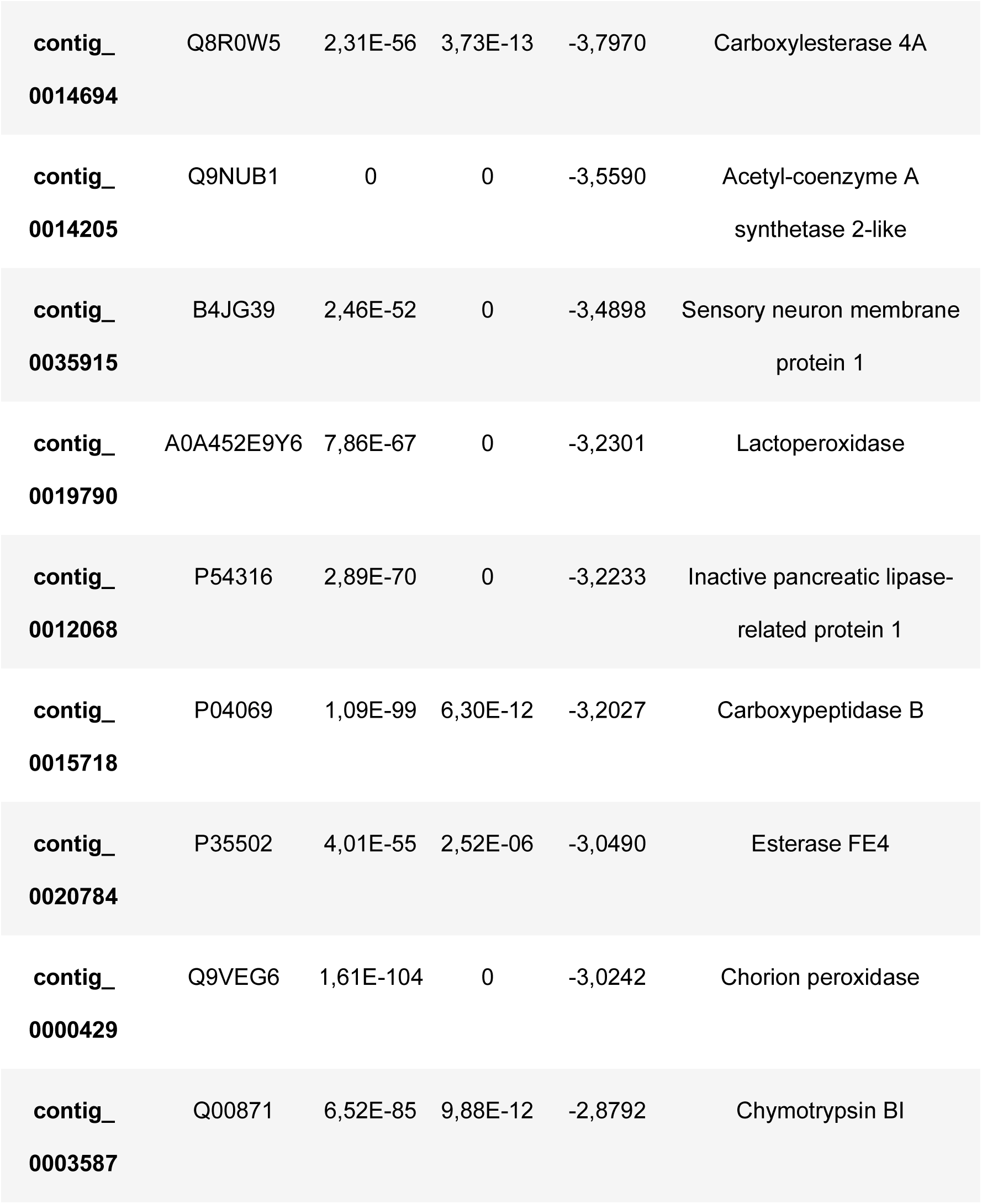
Differentially expressed genes (DEGs) in juvenile *Artemia franciscana* exposed for 24 h to neem extract, compared with untreated controls. The table lists the top 10 upregulated and top 10 downregulated transcripts, ranked by |log₂ fold change|. For each contig, UniProt accession, lowest BLASTx E-value, FDR-adjusted p-value and log₂ fold change are shown together with the corresponding protein annotation.

Several genes encoding enzymes related to metabolism and detoxification showed marked reductions in expression levels. These included *esterase FE4* (log₂FC = −4.24) and *carboxylesterase 4A* (log₂FC = −3.80), both esterases commonly associated with xenobiotic processing, as well as an *acetyl-CoA synthetase 2-like* enzyme (log₂FC = −3.56), involved in acyl-CoA formation. Transcripts annotated as *lactoperoxidase* (log₂FC = −3.23), a heme-containing oxidoreductase, and a *pancreatic lipase-related protein* (log₂FC = −3.22), implicated in lipid hydrolysis, were also significantly downregulated in neem-treated juveniles.

Gene Ontology (GO) enrichment analysis in juveniles (**Fig 3B**) showed significant over-representation of metabolic, immune, and structural functional categories. For biological processes, carbohydrate and polysaccharide catabolic processes were among the most strongly enriched terms. Immune-related categories were also enriched, including terms associated with innate and stress-related immune responses. For molecular function, the term “structural constituent of cuticle” showed the highest enrichment, reflecting consistent changes in genes encoding cuticle-associated proteins.

### 3.5. Targeted analysis of chitin-related transcripts in juvenile A. franciscana

Building on the GO enrichment results, which highlighted cuticle-associated functional categories in juveniles, we quantified transcripts for eight genes associated with chitin biosynthesis and degradation in juvenile *A. franciscana*. Seven of these genes were significantly downregulated in neem-treated juveniles compared with controls. The strongest reductions were observed for *hexose phosphate aminotransferase 1* and *chitin synthase chs2* (**Fig 4A,B**), which encode enzymes involved in precursor production and chitin polymerization, respectively. Overall, neem-treated juveniles showed lower transcript abundance for several chitin-pathway components during early developmental stages.

**Fig 4.**
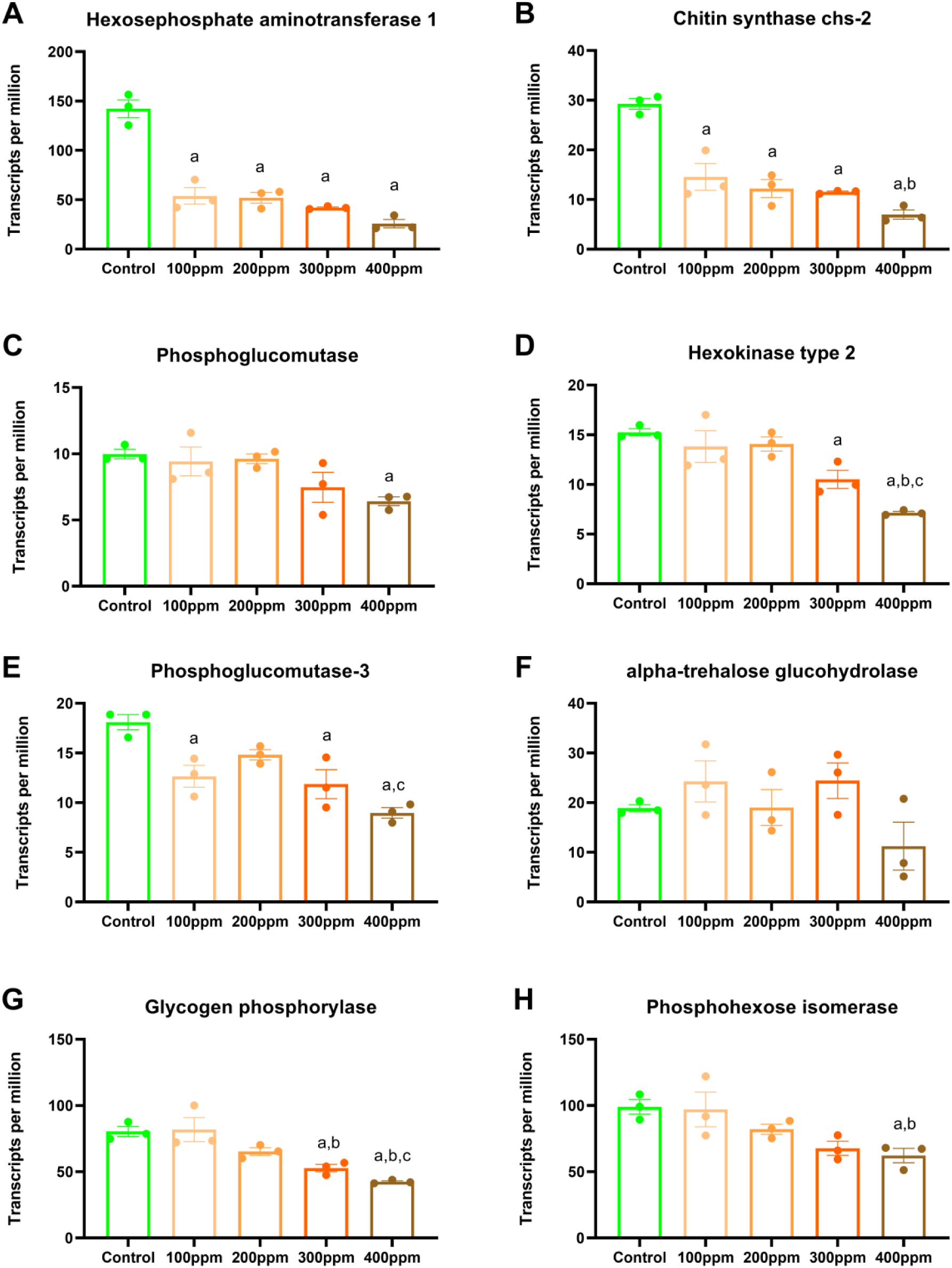
Expression of chitin-related genes in juvenile *A. franciscana* exposed to neem extract. Transcripts per million (TPM) for selected genes involved in chitin precursor production and chitin polymerization in juveniles exposed for 24 h to 0, 100, 200, 300, and 400 ppm neem extract. Panels show (A) hexosephosphate aminotransferase 1, (B) chitin synthase chs-2, (C) phosphoglucomutase, (D) hexokinase type 2, (E) phosphoglucomutase-3, (F) alpha-trehalose glucohydrolase, (G) glycogen phosphorylase, and (H) phosphohexose isomerase. Bars show mean ± SEM (n = 3 libraries per treatment). Different lowercase letters indicate significant differences among treatments (one-way ANOVA followed by Tukey’s multiple comparison test).

### 3.6. Transcript abundance patterns in adult A. franciscana exposed to neem are associated with stress, metabolism, and structural functions

Exposure to neem extract was associated with transcript abundance differences in adult *A. franciscana*. Hierarchical clustering of DEGs showed that untreated samples generally formed a distinct cluster from treated groups (**Fig 5A**). One sample from the 500 ppm treatment (AFATT2_1_1) grouped with controls, indicating within-group variation. Despite this, most treated samples exhibited transcriptional shifts compared to controls. Within the treated groups, adults exposed to 500 ppm displayed a transcriptional profile that differed from both the 250 ppm group and untreated controls.

**Fig 5.**
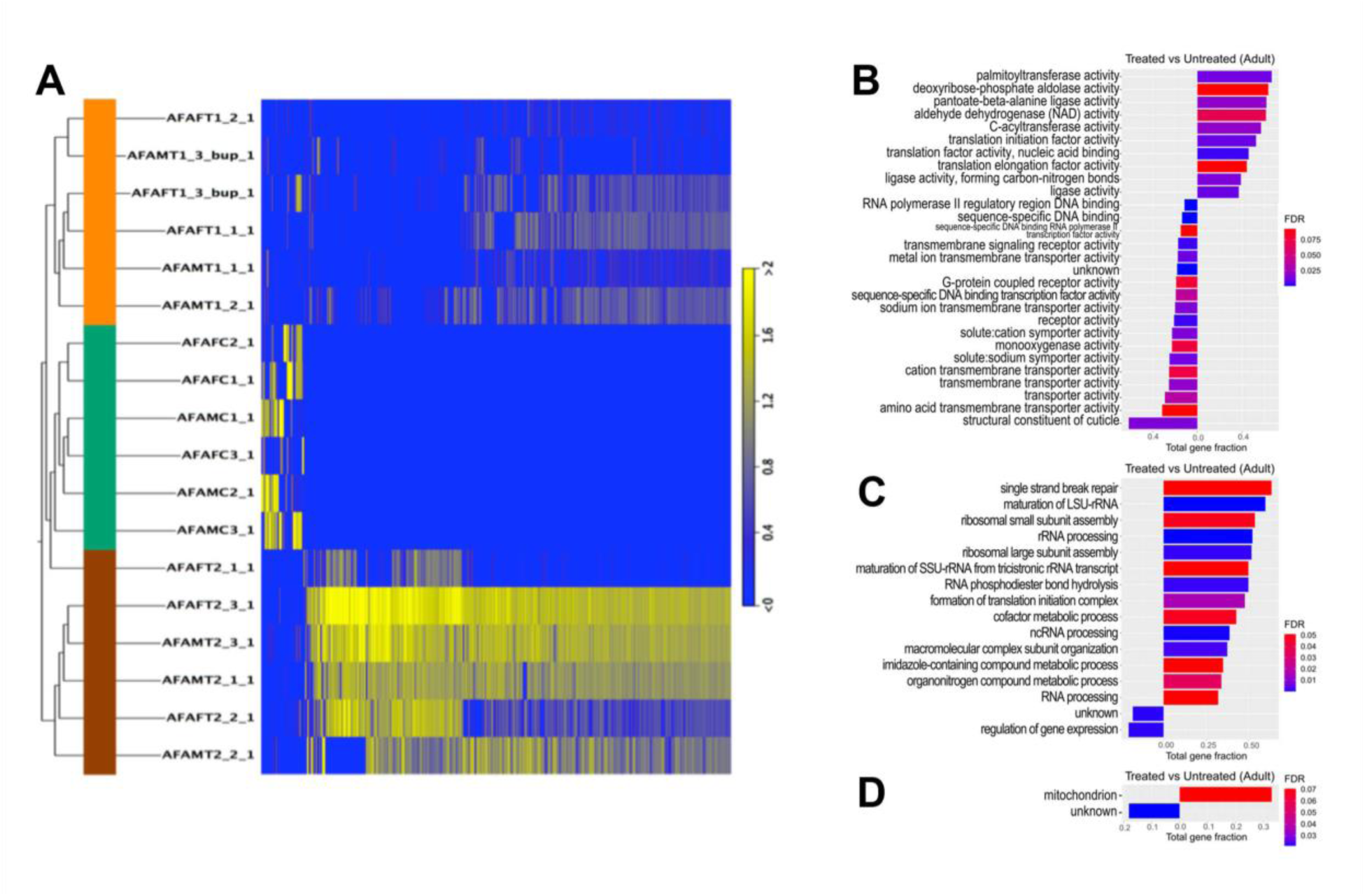
Transcriptomic responses to neem extract exposure in adult *A. franciscana*. **(A)** Heat map of hierarchical clustering of DEGs from adult *A. franciscana* exposed for 7 days to 0 (control), 250 and 500 ppm neem (n = 3 libraries per sex and treatment). Colours indicate relative transcript abundance (low to high). (**B–D)** Enriched Gene Ontology (GO) categories for molecular function **(B)**, biological process **(C)** and cellular component **(D)** in treated (left) and untreated (right) adults. Bar length indicates the number of DEGs assigned to each GO term. “Total gene fraction” denotes the proportion of all annotated genes in the transcriptome associated with that term. Only terms with P < 0.01 are shown. The colour gradient reflects false discovery rate (FDR), with blue indicating lower FDR (higher confidence) and red indicating higher FDR.

Differential expression analysis (**Table 3**) identified several genes with large changes in expression in neem-exposed adults. Strongly upregulated transcripts included *elongation factor 1-alpha* (log₂FC = 14.09), a translation factor involved in protein synthesis, and *tubulin alpha chain* (log₂FC = 13.54), a major component of the cytoskeleton. *Heat shock protein 70 protein 4 (HSPA4)* (log₂FC = 12.07), a molecular chaperone of the HSP70 family, was also highly induced, as was a *major cysteine proteinase* (log₂FC = 13.18), a protease associated with protein turnover.

**Table 3.** Differentially expressed genes (DEGs) in adult *Artemia franciscana* exposed for 7 days to neem extract, compared with untreated controls. The table lists the top 10 upregulated and top 10 downregulated transcripts, ranked by |log₂ fold change|. For each contig, UniProt accession, lowest BLASTx E-value, FDR-adjusted p-value and log₂ fold change are shown together with the corresponding protein annotation.

Upregulated genes
| <u>Name</u> | <u>UniProt</u> | <u>Lowest E-</u> | <u>FDR p-</u> | <u>Log2 fold</u> | <u>Protein name</u> |
| --- | --- | --- | --- | --- | --- |
|  | <u>accession</u> | <u>value</u> | <u>value</u> | <u>change</u> |  |
|  | <u>number</u> |  |  |  |  |
| contig_0005285 | P90519 | 9.95E-116 | 3.54E-11 | 14.0898 | Elongation factor 1-alpha |
| contig_<br>0030322 | P04106 | 0 | 1.19E-10 | 13.5367 | Tubulin alpha chain |
| contig_<br>0036860 | P25779 | 3.73E-113 | 6.10E-10 | 13.1809 | Major cysteine proteinase |
| contig_<br>0017701 | A5DI11 | 0 | 8.40E-09 | 12.4030 | Elongation factor 2 |
| contig_<br>0064049 | P31692 | 3.40E-84 | 1.62E-08 | 12.1311 | Adenine nucleotide<br>translocator |
| contig_<br>0022295 | P11145 | 5.99E-99 | 2.97E-08 | 12.0665 | Heat shock 70 kDa protein<br>4 |
| contig_<br>0026696 | P04106 | 0 | 8.02E-08 | 11.9981 | Tubulin alpha chain |
| contig_<br>0071908 | P26796 | 3.27E-122 | 3.23E-08 | 11.8823 | 60S acidic ribosomal<br>protein P0 |
| contig_<br>0043702 | O13672 | 2.50E-60 | 4.99E-08 | 11.8531 | 60S ribosomal protein L8 |
| contig_<br>0039289 | P69104 | 1.16E-122 | 4.75E-08 | 11.8479 | Guanine nucleotide-binding<br>protein subunit beta-like<br>protein |

Downregulated genes
| <u>Name</u> | <u>UniProt</u> | <u>Lowest E-</u> | <u>FDR p-</u> | <u>Log2 fold</u> | <u>Protein name</u> |
| --- | --- | --- | --- | --- | --- |
|  | <u>accession</u> | <u>value</u> | <u>value</u> | <u>change</u> |  |
|  | <u>number</u> |  |  |  |  |
| contig_<br>0055157 | Q9JME2 | 7.25E-61 | 1.34E-08 | -6.6510 | Carbohydrate<br>sulfotransferase 11 |
| contig_<br>0086110 | P13706 | 1.46E-53 | 0.00717 | -4.8626 | Glycerol-3-phosphate<br>dehydrogenase [NAD(+)],<br>cytoplasmic |
| contig_<br>0022053 | P00765 | 5.47E-51 | 4.02E-05 | -3.0847 | Trypsin-1 |
| contig_<br>0005802 | P00771 | 2.96E-53 | 0.00063 | -2.9908 | Collagenolytic protease |
| contig_<br>0032565 | Q9S777 | 7.65E-65 | 9.71E-05 | -2.9622 | 4-coumarate--CoA ligase 3 |
| contig_<br>0002900 | P09470 | 8.25E-82 | 0.00512 | -2.7686 | Angiotensin-converting<br>enzyme |
| contig_<br>0015718 | P04069 | 1.09E-99 | 2.20E-05 | -2.7198 | Carboxypeptidase B |
| contig_<br>0014694 | Q8R0W5 | 2.31E-56 | 8.90E-08 | -2.6582 | Carboxylesterase 4A |
| contig_<br>0057631 | P27619 | 2.12E-98 | 0.03036 | -2.6305 | Dynamin |
| contig_<br>0000432 | Q04960 | 2.13E-95 | 0.00055 | -2.5258 | DnaJ protein homolog |

Several genes associated with energy metabolism and other homeostatic functions showed reduced expression levels in adults exposed to neem. *Glycerol-3-phosphate dehydrogenase*, an enzyme involved in glycolysis and lipid metabolism, was strongly downregulated (log₂FC = −4.86), as was *carbohydrate sulfotransferase 11* (log₂FC = −6.65), which participates in glycosaminoglycan biosynthesis.

Gene Ontology (GO) enrichment analysis in adults (**Fig 5B–D**) showed significant over-representation of categories related to “regulation of gene expression” and “transcription factor activity” in treated samples. In contrast, untreated adults exhibited higher representation of GO terms such as “ribosomal large subunit assembly” and “rRNA processing”, as well as lipid metabolic processes including lipase and aldehyde dehydrogenase activities. Treated adults also showed enrichment of structural molecule activity associated with the extracellular region and cuticle.

Collectively, the gene-expression and GO patterns in adults highlight stress-, metabolism-and cuticle-associated responses to neem exposure, supporting a closer examination of chitin-related transcripts.

### 3.7. Chitin-related gene expression in adult A. franciscana exposed to neem

To determine whether chitin-associated pathways were also affected in adults, we analysed the same panel of eight genes related to chitin biosynthesis and degradation used for juveniles in neem-treated and untreated adult *A. franciscana* (**Fig 6**). Three of these genes showed significant expression differences between treatments. In males, *hexokinase type 2* and *phosphohexose isomerase*, both associated with carbohydrate metabolism and chitin precursor synthesis, were significantly downregulated in neem-exposed individuals compared with controls (**Fig 6A,B**). In females, *glycogen phosphorylase*, an enzyme contributing to mobilization of stored carbohydrates toward biosynthetic pathways, showed reduced expression in treated samples (**Fig 6C**).

**Fig 6.**
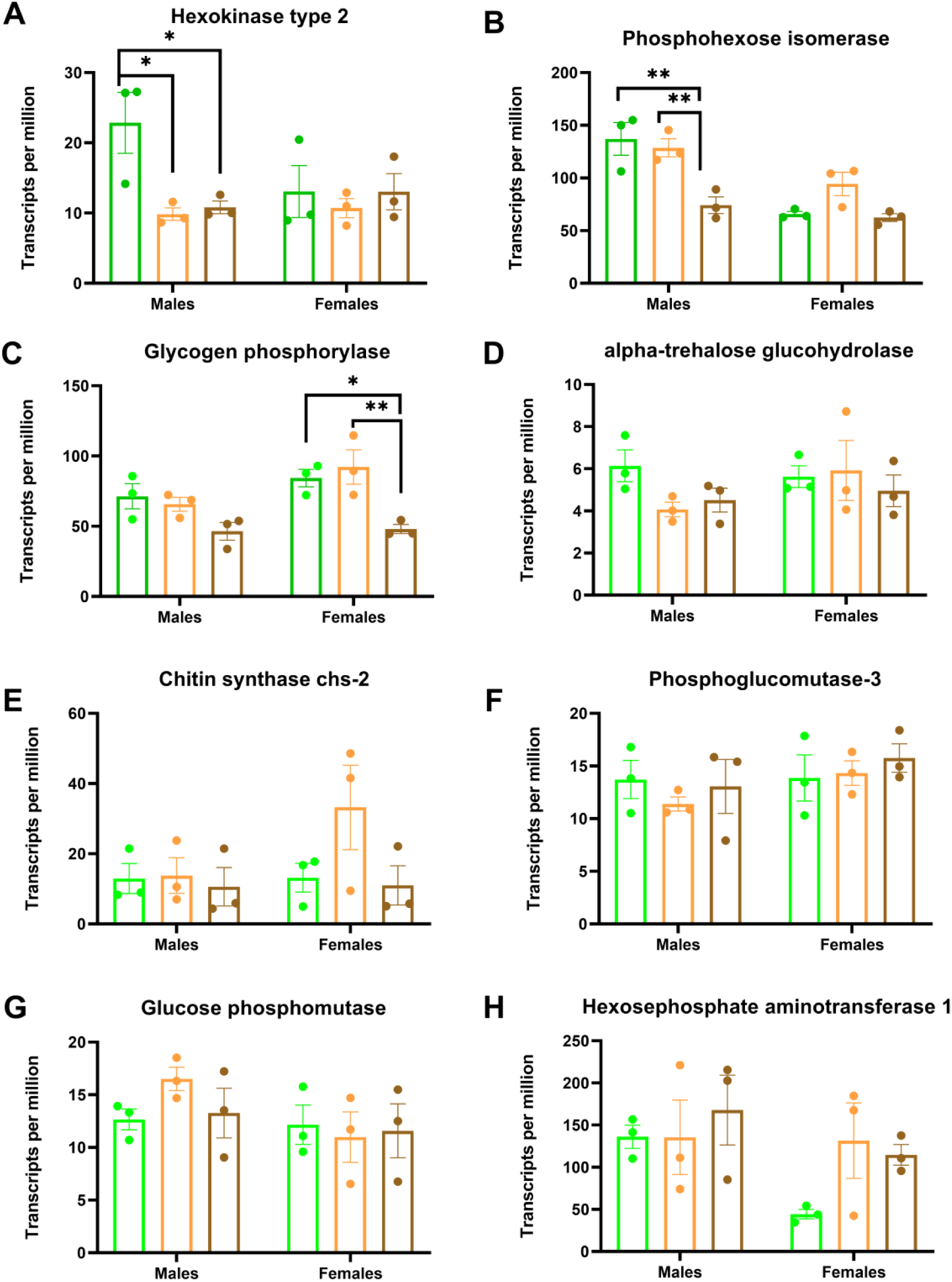
Expression of chitin-related genes in adult *A. franciscana* exposed to neem extract. Transcripts per million (TPM) for selected genes involved in chitin precursor metabolism in adult males and females exposed for 7 days to 0, 250, and 500 ppm neem extract. Panels show (A) hexokinase type 2, (B) phosphohexose isomerase, (C) glycogen phosphorylase, (D) alpha-trehalose glucohydrolase, (E) chitin synthase chs-2, (F) phosphoglucomutase-3, (G) glucose phosphomutase, and (H) hexosephosphate aminotransferase 1. Bars show mean ± SEM (n = 3 libraries per sex and treatment). Brackets indicate the pairwise comparisons tested, and asterisks indicate statistically significant differences between groups (*P < 0.05, **P < 0.01; two-way ANOVA followed by Tukey’s multiple comparison test).

### 3.8. Histological analysis of adult Artemia

Histological analysis revealed progressive structural alterations in the embryonic brood sac (EBS) cuticle of females exposed to neem extract. In untreated control samples (0 ppm; **Fig 7A-C**), the EBS displayed a smooth, well-defined cuticle (arrowheads) enclosing the embryos. The cuticle appeared uniformly thick, with intact structural integrity. The moon-shaped embryos within the brood sac corresponded to desiccating *A. franciscana* cysts (arrows) and exhibited normal morphology. At 100 ppm and 250 ppm neem extract (**Fig 7D-I**), the cuticle showed irregularities compared to controls, including unevenness in shape, variations in thickness, and a less well-defined morphology. More pronounced deformities were observed at 500 ppm (**Fig 7J-L**), where the EBS cuticle exhibited severe thinning and fragmentation, with some regions appearing poorly developed or disrupted. Additionally, the desiccating embryos (arrows) showed irregular morphology compared to controls.

**Fig 7.**
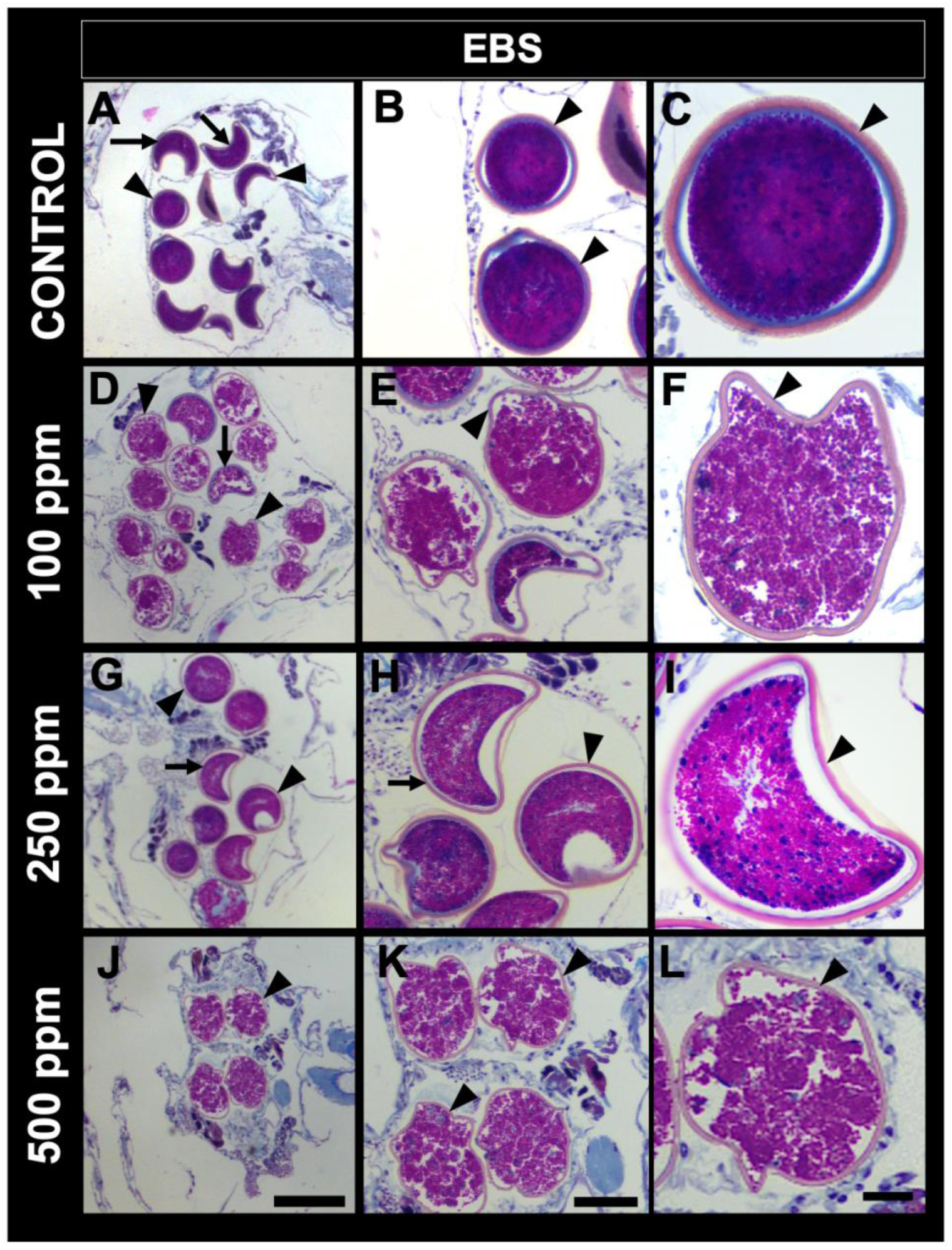
Detail of the cuticle enclosing the embryos in the brood sacs (EBS) in the adult female reproductive tract of *Artemia franciscana*. Arrowheads: cuticle. Arrow: Desiccating Artemia cysts. (**A-C**): Untreated control sample. Smooth cuticle enclosing the EBS. (**D-F**): Neem treatment with 100 ppm. Irregular EBS cuticle morphology is shown. (**G-I**) Neem treatment with 250 ppm. (**J-L**): Neem treatment with 500 ppm. Irregular EBS cuticle morphology is shown. Moon-shaped embryos enclosed with cuticle inside the brood sacs correspond to *Artemia* cysts in the process of desiccation. (A, D, G, J): Bar = 200 µm. (B, E, H, K): Bar = 100 µm. (C, F, I, L): Bar = 50 µm.

Beyond the cuticle, neem exposure impacted the internal reproductive organs of *Artemia*. Morphometric analysis showed that females from higher neem treatments had a significantly reduced ovarian area (P = 0.0006) compared to controls (**S1 Fig**). In healthy brine shrimp, the ovary is a prominent organ where oocytes mature, but in neem-treated females ovarian area was reduced. In contrast, males did not show significant changes in testis size.

## 4. Discussion

Neem exposure produced strong, dose-dependent effects in *A. franciscana*, affecting both juveniles and adults. Larvae exhibited high mortality and abnormal phenotypes at concentrations ≥300 ppm, while adults displayed significant mortality after 7-day exposures to 500 ppm. Males were more susceptible than females, a pattern consistent with documented sex-specific sensitivities to toxicants in crustaceans (26–29). Baseline transcriptomic studies have shown that males and females of *A. franciscana* differ markedly in gene expression profiles (30,31). Together with our transcriptomic and histological data, these phenotypic outcomes indicate that neem-derived compounds were associated with marked changes in survival, development, and reproductive-associated endpoints in this model crustacean.

Transcriptomic analysis suggested life stage- and sex-specific transcript abundance patterns associated with neem exposure. While both juveniles and adults mounted transcriptomic responses, adults exhibited more extensive changes (19,957 DEGs vs. 933 in juveniles), suggesting heightened transcriptional reprogramming. Such widespread changes reflect the activation of stress response pathways and suppression of metabolic processes, consistent with toxicant-induced physiological disruption in other aquatic arthropods (32–34). A comparable pattern has been described in insect larvae exposed to azadirachtin, where large-scale transcriptomic reprogramming affected more than 1,200 genes in *Spodoptera frugiperda*, including detoxification and structural genes, together with suppression of chitin synthase and other chitin- and cuticle-associated genes involved in cuticle formation (35). In line with these findings, one of the most prominent transcriptomic signatures in juvenile *A. franciscana* was the disruption of chitin metabolism, with seven of eight analyzed chitin-related genes significantly downregulated, including *hexose phosphate aminotransferase* and *chs2*, which are critical for chitin precursor production and polymerization during exoskeleton formation. Adults also exhibited significant transcriptomic disruptions, particularly upregulation of heat shock proteins (e.g., HSPA4), tubulin, and cysteine proteases, indicating activation of stress, cytoskeletal reorganization, and tissue remodeling responses. Downregulated transcripts were enriched for energy metabolism and protein biosynthesis functions, including glycolytic enzymes, sulfotransferases, and ribosomal proteins. These patterns suggest a shift from growth-related processes to cellular maintenance and survival—a known stress response in arthropods (36,37).

Expression of chitin-related genes in adults showed selective downregulation of enzymes involved in carbohydrate metabolism, notably in males. Reduced expression of *hexokinase type 2* and *phosphohexose isomerase* in males, and *glycogen phosphorylase* in females, is consistent with reduced capacity for UDP-N-acetylglucosamine production, the key precursor for chitin biosynthesis. These enzymes participate in the early steps of the hexosamine pathway supplying substrates for chitin formation (38–40), and glycogen phosphorylase is known to mobilize stored carbohydrates into this biosynthetic route (39,41). Since chitin synthesis underpins the structural integrity of crustacean exoskeletal and reproductive tissues, including brood sac and embryonic envelopes (42,43), we speculate that these molecular alterations may contribute to the brood sac changes observed histologically.

Histopathological analysis provided morphological evidence of treatment-associated effects. Treated females exhibited structural alterations of the embryonic brood sac cuticle and significantly reduced ovarian areas. These observations are consistent with the transcript abundance patterns identified in the transcriptomic analysis, particularly those involving cuticle-associated and chitin-related genes. Similar reproductive impairments have been reported in other invertebrates exposed to neem-based products. In *Daphnia*, chronic neem exposure reduced offspring viability (44), while in *Bombus terrestris*, azadirachtin ingestion caused ovary atrophy and cessation of egg-laying (45).

From a formulation and chemistry perspective, it is also important to note the compositional distinctions between neem kernel and neem oil. Neem kernel is rich in azadirachtin and related limonoids, while neem oil contains primarily fatty acids and acts as a delivery enhancer. The extract used in our study derived from kernel content, where azadirachtin is a dominant component known to inhibit chitin synthesis and disrupt molting (46,47). Although this was not directly tested here, the potential synergy between oil-based lipids and kernel-derived limonoids may amplify toxicological outcomes in certain contexts (48,49). Understanding these chemical distinctions is essential for interpreting variability across studies and guiding safe application in aquaculture settings.

Although neem is often perceived as an environmentally benign biopesticide, several studies have shown that neem-based formulations can exert lethal and sublethal effects on non-target aquatic organisms, including invertebrates and fish (7,44,50,51). Our results add to this evidence by showing that, under the laboratory conditions used here, neem exposure was associated with phenotypic abnormalities, histological alterations, and transcriptomic changes in *A. franciscana*. Given that zooplanktonic crustaceans play central roles in many aquatic food webs, comparable sublethal effects in ecologically relevant taxa could, in principle, translate into altered population dynamics and ecosystem functioning (16,17). The combined use of transcriptomic and histological endpoints in the present study illustrates how mechanistic biomarkers can reveal early signs of stress and toxicity before overt population-level declines become apparent and supports their broader application in ecotoxicological risk assessment (8,12,52).

Finally, this study highlights the utility of combining high-throughput transcriptomics with classical ecotoxicological endpoints. Gene expression profiling provided molecular information on stress, detoxification, and structural impairments that are not discernible from phenotypic observations alone, consistent with the growing use of omics-based tools in environmental monitoring and pesticide risk assessment (8,9). In summary, neem extract was associated with changes in *A. franciscana* related to exoskeletal maintenance, metabolism, and reproduction, together with transcriptomic and histological alterations. These observations are relevant to ongoing efforts to integrate molecular data into structured ecotoxicological interpretation frameworks, including Adverse Outcome Pathway–based approaches (53). Together, these results contribute to understanding of how botanical insecticides affect non-target aquatic invertebrates and support careful evaluation of their use in habitats supporting sensitive species.

## Acknowledgements

Donald Brown contributed significantly to this work and is included as a co-author posthumously. We dedicate this paper to his memory.

## Data Availability Statement

All raw sequencing data generated in this study are publicly available in the NCBI Sequence Read Archive (SRA) under BioProject accession PRJNA1402122. The de novo assembled *Artemia franciscana* transcriptome is provided in S1 Dataset, the BLAST-based annotation table is provided in S2 Dataset, the juvenile transcript-abundance dataset is provided in S3 Dataset, and the adult transcript-abundance dataset is provided in S4 Dataset. The cDNA library metadata are provided in S1 Table. All other data underlying the findings are provided within the manuscript and its Supporting Information files.

## Supporting information

**S1 Table.** cDNA libraries included in the RNA-seq experiment for juvenile and adult *Artemia franciscana*. For each library, sample name, developmental stage, sex, treatment condition, and neem concentration are indicated.

**S1 Dataset.** De novo assembled Artemia franciscana transcriptome in FASTA format.

**S2 Dataset.** BLAST-based annotation table for the de novo assembled Artemia franciscana transcriptome.

**S3 Dataset.** Juvenile expression browser containing transcript abundance data for Artemia franciscana juveniles exposed to neem extract.

**S4 Dataset.** Adult expression browser containing transcript abundance data for Artemia franciscana adults exposed to neem extract.

**S1 Video.** Representative stereomicroscope recordings of juvenile (naupliar-stage) Artemia franciscana exposed for 24 h to neem extract. On-screen labels are shown in g/L and correspond to 0 (control), 100, 200, 300, 400, 500, 600, 700, and 800 mg/L. Normal juveniles show active swimming with coordinated, undulating movements of the podia and maintain their position in the water column. Abnormal juveniles display reduced and/or uncoordinated podial movements and are unable to maintain their position in the water column. Dead juveniles remain immobile on the well bottom, show no response to gentle tapping, and may exhibit necrotic morphology.

**S2 Video.** Reference video for adult sex identification in control Artemia franciscana. Representative stereomicroscope recordings of unexposed adult A. franciscana of both sexes used for the 7-day survival experiment. Males are identified by the presence of claspers, and females are shown as gravid (ovoviviparous).

**S1 Fig.** Gonadal area of adult Artemia franciscana exposed to neem extract. Gonadal area of females and males after 7 days of exposure to 0, 100, 250, and 500 mg/L neem extract. (A) Ovarian area in females. (B) Testis area in males. Data are shown as mean ± SD (n = 10 individuals per sex and treatment). Asterisks indicate significant differences from the corresponding control group (one-way ANOVA followed by Tukey’s multiple comparison test, P < 0.05); no significant differences were detected among male treatments.

## Notes

### Competing Interest Statement

The authors have read the journal?s policy and have the following competing interests: A.E. and P.P. are employees of Cargill, Inc. J.P. was affiliated with Cargill, Inc. during the conduct of the study. Their specific roles are articulated in the Author Contributions section. This does not alter our adherence to PLOS ONE policies on sharing data and materials. The authors declare that no other competing interests exist.

